# *Plasmodium knowlesi* clinical isolates from Malaysia show extensive diversity and strong differential selection pressure at the merozoite surface protein 7D (MSP7D)

**DOI:** 10.1101/537621

**Authors:** Md Atique Ahmed, Fu-Shi Quan

## Abstract

**Background:** High proportion of human cases due to the simian malaria parasite *Plasmodium knowlesi* in Malaysia has been a cause of concern, as it can be severe and fatal. Merozoite surface protein 7 (MSP7) is a multigene family which forms a non-covalent complex with MSP-1 prior to receptor-ligand recognition in *Plasmodium falciparum* and thus an important antigen for vaccine development. However, no study has been done in any of the ortholog family members in *Plasmodium knowlesi* from clinical samples. Thus in this study we investigated the level of polymorphism, haplotypes, and natural selection acting at the *pkmsp-7D* gene in clinical samples from Malaysia.

**Methods:** We analyzed 36 full-length *pkmsp7D* gene sequences (along with the reference H-strain: PKNH_1266000) which were orthologous to *pvmsp7H* (PVX_082680) from clinical isolates of Malaysia available from public databases. Population genetic, evolutionary and phylogenetic analyses were performed to determine the level of genetic diversity, polymorphism, recombination and natural selection.

**Results:** Analysis of 36 full-length *pkmsp7D* sequences identified 147 SNPs (91 non-synonymous and 56 synonymous substitutions). Nucleotide diversity across the full-length gene was higher than its ortholog in *P. vivax* (*msp7H*). Region-wise analysis of the gene indicated that the nucleotide diversity at the central region was very high (π = 0.14) compared to the 5’ and 3’ regions. Most hyper-variable SNPs were detected at the central domain. Multiple test for natural selection indicated the central region was under strong positive natural selection however, the 5’ and 3’ regions were under negative/purifying selection. Evidence of intragenic recombination were detected at the central region of the gene. Phylogenetic analysis using full-length *msp7D* genes indicated there was no geographical clustering of parasite population.

**Conclusions:** High genetic diversity with hyper-variable SNPs and strong evidence of positive natural selection at the central region of MSP7D indicated exposure of the region to host immune pressure. Negative selection at the 5’ and the 3’ regions of MSP7D might be because of functional constraints at the unexposed regions during the merozoite invasion process of *P. knowlesi*. No evidence of geographical clustering among the clinical isolates from Malaysia indicated uniform selection pressure in all populations. These findings highlight the further evaluation of the regions and functional characterization of the protein as a potential blood stage vaccine candidate for *P. knowlesi*.

## Background

Malaria is a major global health problem with high mortality and morbidity rates and thus a significant barrier to socio-economic development of under-developed countries [1]. Among all the malarias, *Plasmodium knowlesi* is a simian malaria parasite which is now considered as the fifth *Plasmodium* species infecting humans and an emerging infection in Southeast Asian countries [2–6]. Countries in Central Asia and almost all Southeast Asia countries has reported human infections due to *P. knowlesi* including Malaysia [4, 7, 8], Singapore [9], Myanmar [10], Vietnam [11], Indonesia [12, 13], Philippines [14], Cambodia [15], India [16] and Thailand [17]. Malaysia being the epicenter of *knowlesi* malaria, infection accounts up to 90% human infections [4, 8, 18, 19]. Despite the overall reduction in malaria cases around the world, the increasing trend of *P. knowlesi* cases in Southeast Asia highlights the need for proper and effective elimnation measures as well as development of effective vaccines.

*P. knowlesi* has a 24-hour erythrocytic cycle thus rapid increase in parasitaemia has been associated with the development of severe malaria and also a common cause for severe and fatal malaria in Malaysian Borneo [3, 20–22]. Studies on mitochondrial and ssrRNA genes in *P. knowlesi* from patients and wild macaques identified two distinct sub-populations which clustered geographically to Peninsular Malaysia and Malaysian Borneo [23]. Additionally, genetic and genomic studies from Malaysia identified 3 distinct sub-populations; two originating from Sarawak and one from Peninsular Malaysia [24–27] thereby highlighting the complexity of infections in humans and the challenges for control as well as vaccine design.

Development of an effective malaria blood-stage vaccine antigens have focused on selecting antigens that provides long term and strain transcending protection and blocks invasion of red blood cells (RBCs). Studies have shown that parasite antigens which are recognized by host’s immune system, accumulate polymorphism as a means of evading host’s defense mechanism and are prime targets for vaccine development. These are achieved through the knowledge of antigen natural selection and the pattern of diversity across geographical areas. Blood-stage antigens in *P. falciparum* (e.g. merozoite surface proteins (MSPs), apical membrane antigen 1 (AMA1) that are under strong positive balancing selection are important targets of such acquired immunity and are supported by antibody inhibition assays in culture as well as naturally acquired antibodies in endemic regions [28, 29]. Several known blood stage antigens (e.g. MSP1, MSP1P and AMA1) have been recently studied in *P. knowlesi*, but most of the antigens were found to be under the influence of strong negative/purifying selection indicating no role in immune evasion mechanism [30–33]. PfMSP1 has been a major target for anti-malaria vaccine development but recent studies have shown that MSP 6 and 7 forms a non-covalent complex with MSP1 during the invasion process [34]. Disruption of PfMSP7 gene has resulted in partial inhibition of erythrocyte invasion during merozoite stages thus indicating that it is an important molecule [35]. Furthermore, MSP7 binding to P-selectin receptor has suggested its role in modulating disease severity in mice models [36] thereby indicating that immunity induced by MSP7 could potentially disrupt parasite development.

MSP7 is a multi-gene family and the number of paralog members in each species differs [37]. For example, till date 13 MSP7 members were identified in *P. vivax* which are arranged in head to tail arrangement in chromosome 12 and are named alphabetically as MSP7A to MSP7M [37]. Three of its members i.e. PvMSP7C, PvMSP7H and PvMSP7I have been found to be highly polymorphic at the central region and under strong positive natural selection in Colombian population indicating it is an important molecule for consideration as a vaccine candidate [38]. *P. falciparum* has 8 MSP7 genes and low genetic polymorphism has been observed within the field isolates [39]. Certain MSP7 members forms part of the protein complex interacting with host cells [40] and they are localized at the merozoite surface [41, 42]. Despite the fact that among the human malarias, *P. knowlesi* is phylogenetically closely related to *P. vivax* and there are five paralog members in *pkmsp7* gene family [37], no genetic study has been done to characterize the *P. knowlesi* MSP7 ortholog members from clinical samples. Thus in this study *pkmsp-7D* gene which is an ortholog to the *Pvmsp-7H* was chosen for genetic analysis.

In this study firstly, 36 *pkmsp-7D* full-length sequences (32 clinical isolates and 4 laboratory lines of Malaysia) were obtained from published databases based on the ortholog gene sequence in *P. vivax msp-7H* (PVX_082680) [37, 43]. The level of sequence diversity, natural selection using full-length genes as well at each of the three regions of the gene (5’, 3’ and the central region) were determined (along with the H-strain) of Malaysia. The information obtained from this study will be helpful for designing functional studies and future rational design of a vaccine against *P. knowlesi*.

## Methods

### *pkmsp-7D* sequence data

Based on the identification of ortholog members in *Plasmodium* species [37] the *pkmsp-7D* sequences were downloaded for 36 isolates (32 clinical and 4 long term isolated lines) originating from Kapit, Betong and Sarikei in Malaysian Borneo and Peninsular Malaysia along with the H-strain (PKNH_1266000, old ID; PK13_3510c) (Additional file 1) [24]. The sequence data with accession numbers are given in (Additional file 1). Signal peptide for the full-length *pkmsp-7D* was predicted using Signal IP 3.0 and the trans membrane regions using the Phobious prediction software [44, 45]. The PkMSP7D regions were characterized based on the published ortholog of PvMSP7H (PVX_082680).

### Sequence diversity and natural selection

Sequence diversity (π), defined as the average number of nucleotide differences per site between two sequences within the sequences, was determined by DnaSP v5.10 software [46]. Number of polymorphic sites, number of synonymous and non-synonymous substitutions, haplotype diversity (Hd), number of haplotypes (h) within the *pkmsp7D* sequences were also determined by DnaSP software.

To investigate departure from neutrality, Tajima’s D analysis was conducted [47]. Under neutrality, Tajimas D is expected to be 0. Significantly, positive Tajima’s D values indicate recent population bottleneck or balancing selection, whereas negative values suggest population expansion or negative selection. Natural selection was estimated using the modified Nei-Gojobori method to calculate the average number of non-synonymous (d_N_) and synonymous (d_S_) substitutions. Difference between d_N_ and d_S_ were determined by applying codon based Z-test (*P* < 0.05) in MEGA software v.5 with 1000 bootstrap replications [48]. Additionally, to test for natural selection in inter-species level, the robust McDonald and Kreitman (MK) test was also performed with closest *msp7*orthologs in both *Plasmodium vivax* (PVX_082680) and *Plasmodium cynomolgi* (PCYB_122830) as outgroups individually using DnaSP v5.10 software.

### Analysis of recombination and linkage disequilibrium

Analysis of the minimum number of recombination events (Rm) within the *pkmsp7D* genes was performed using DnaSP software v5.10 software [46]. Linkage disequilibrium (LD) is the non-random association of sequences at two or more loci. Linkage disequilibrium index, r^2^ (square of the correlation coefficient of allelic states at each pair of loci) against nucleotide distance were plotted for *pkmsp7D* sequences across the gene using DnaSP 5.10 software. A statistical significance test was conducted using Fisher's exact test and □^2^ test departures from randomness and the value of r^2^ ranges from 0 to 1 in the software.

### Phylogenetic analysis

Phylogenetic analysis was conducted using deduced amino acid sequences from 32 PkMSP7 full-length sequences from Malaysian Borneo, 5 laboratory lines from Peninsular Malaysia; reference H-strain (PKNH_1266000), the Malayan Strain (PKNOH_S09532900), MR4H strain, Philippine strain and the Hackeri strain along with other *P. knowlesi* MSP7 paralog members PkMSP7A (PKNH_1266300), PkMSP7E (PKNH_1265900) and PkMSP7C (PKNH_1266100) were included. Other ortholog members of *Plasmodium vivax* MSP7H (PVX_082680), MSP7E (PVX_082665) and *Plasmodium coatneyi* (PCOAH_00042440) were also included for comparative analysis. Maximum Likelihood (ML) method based on Poisson correction model was used as described in MEGA 5.0 with 1,000 bootstrap replicates to test the robustness of the trees.

## Results

### Polymorphism within full-length *pkmsp7D* genes and natural selection

The Signal IP server detected a signal peptide in between amino acid positions 22 and 23 of the PkMSP7D protein (Additional file 2) however, no transmembrane region was detected (data not shown). Alignment and comparison of the amino acid sequences of the full-length *P. knowlesi* H reference strain MSP7D sequence with *P. vivax* MSP7H Sal-1 (PVX_082680) reference strain showed 60.4% identity and the next nearest PvMSP7 ortholog gene was *E* which had 47.3% identity. The amino acid identity within the PkMSP7 paralog members (A, C, D and E) were in the range of 17-26%. The schematic structure of PkMSP7D protein gene with demarcated regions are shown in Fig. 1. The two cysteine residues at the 5’ and 3’ regions were found to be conserved all the 36 sequences. Within the full-length *pkmsp7D* sequences (n = 36), there were 203 (17.13%) polymorphic sites leading to 34 haplotypes (Table 1). Of the 203 SNPs, 157 were parsimony informative sites, 46 were singleton variable sites. Among the 157 parsimony informative sites, 131 were two variant, 23 were three variants and 3 were four variants. Parsimony informative sites with 3 or 4 variants led to highly variable non-synomymous substitutions within the central region (Fig. 2). Overall, within the full-length gene, there were 147 SNPs which could be analyzed by DnaSP (91 non-synonymous substitutions and 56 synonymous substitutions) (Table 2) and 75 non-synonymous complex codons (which were highly variable within the central region) were excluded from analysis by the DnaSP software. The overall nucleotide diversity was π = 0.052 ± SD 0.002 which was higher than its ortholog in *P. vivax msp7H* [38] and the haplotype diversity was 0.998 ± SD 0.002 (Table 1). Sliding window analysis of diversity indicated that highest diversity was at the central region of the gene (Fig. 3A). The sliding window analysis of Tajimas’D indicated the central region of the gene had the highest D values (Fig. 3B). Interestingly, the 3’ region also had some SNPs, which had positive D values (Fig. 3B). The natural selection analysis of the full-length *pkmsp7D* genes indicated d_N_-d_S_ = −2.1, however, Tajimas’ D and Fu and Li’s D* and F* values were positive (Table 1). The robust inter-species McDonald-Kreitman test indicated a positive selection when *P. vivax* (NI = 1.25, *P > 0.1*) and *P. cynomolgi* (NI = 1.38, *P > 0.1*) were used as out-groups but were not significant (Table 3). Nucleotide polymorphisms across the full-length gene with hyper-variable SNPs within the sequences are shown in Additional file 3.

**Table 1.** Estimates of nucleotide diversity, natural selection, haplotype diversity and neutrality indices of *pkmsp7*

| Region | No.<br>samples | SNPs | No.<br>haplotype | Diversity ± SD | | $d_N-d_S$ | Codon<br>based $z$<br>test | Taj D | Fu &<br>Li's D* | Fu &<br>Li's F* |
| --- | --- | --- | --- | --- | --- | --- | --- | --- | --- | --- |
|  |  |  |  | Haplotype | Nucleotide |  |  |  |  |  |
| Full-length | 36 | 203 | 34 | $0.998 \pm 0.007$ | $0.052 \pm 0.002$ | -2.0 | $P > 0.1$ | 0.295 | 0.190 | 0.270 |
| 5' region | | 28 | 30 | $0.992 \pm 0.009$ | $0.010 \pm 0.001$ | -3.0 | $P > 0.1$ | -1.25 | -2.24 | -2.26 |
| Central<br>region | | 151 | 32 | $0.995 \pm 0.008$ | $0.1495 \pm 0.006$ | 1.84 | $P < 0.05$ | 0.673 | 0.896 | 0.973 |
| 3' region | | 32 | 30 | $0.990 \pm 0.010$ | $0.017 \pm 0.0007$ | -2.68 | $P < 0.01$ | -0.275 | -1.027 | -0.917 |

*Abbreviations:* SNPs single nucleotide polymorphisms; SD Standard deviation

**Table 2.** Synonymous and non-synonymous sites of *pkmsp7*

| Location | n | SNPs | Syn. | Non-syn. |
| --- | --- | --- | --- | --- |
| Full-length |  | 147 | 56 | 91 |
| 5' region |  | 28 | 18 | 10 |
| Central region | 36 | 89 | 16 | 73 |
| 3' region |  | 30 | 22 | 8 |

*Abbreviations:* SNPs Single nucleotide polymorphisms; Syn. Synonymous substitutions; Non-syn. Nonsynonymous substitutions; n number of samples

**Table 3.** McDonald–Kreitman tests on MSP7D of *Plasmodium knowlesi* and its regions with *P.vivax* and *P. cynomolgi* orthologs as outgroup species.

| MSP7 | Polymorphic changes within *P. knowlesi* |  | Fixed differences between species |  |  |  | Neutrality Index |  |
| --- | --- | --- | --- | --- | --- | --- | --- | --- |
|  | Syn | NonSyn | *Pk* vs *Pv*<br>Syn | *Pk* vs *Pv*<br>NonSyn | *Pk* vs *Pcy*<br>Syn | *Pk* vs *Pcy*<br>NonSyn | *Pk* vs *Pv* | *Pk* vs *Pcy* |
| Full-length | 56 | 91 | 145 | 188 | 112 | 132 | 1.25 | 1.38 |
| 5' region | 18 | 10 | 47 | 51 | 52 | 51 | 0.51 | 0.56 |
| Central region | 16 | 73 | 31 | 57 | 15 | 30 | 2.49* | 2.28* |
| 3' region | 22 | 8 | 67 | 78 | 54 | 61 | 0.31<sup>‡</sup> | 0.32<sup>‡</sup> |

Abbreviations: \* Fisher's exact test P-value < 0.05; <sup>‡</sup> Fisher's exact test P-value < 0.01; Syn Synonymous sites; NonSyn Non synonymous sites; *Pk* *Plasmodium knowlesi*; *P.v* *Plasmodium vivax*; *P.cy* *Plasmodium cynomolgi*

**Fig. 1.**
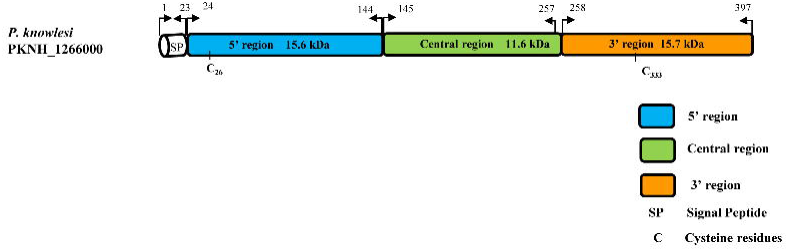
Schematic diagram of PKMSP7 protein based on the H-Strain (PKNH_1266000). Each box in the schematic diagram is representative of the regions found in PkMSP7 and their respective amino acid positions. The central region is represented in green and blue and orange region represents the 5’ and the 3’ regions respectively. The two 6-Cys residues found within the protein are marked along with approximate molecular weight of each region. Signal peptide is abbreviated as (SP) and cysteine residues as C.

**Fig. 2.**
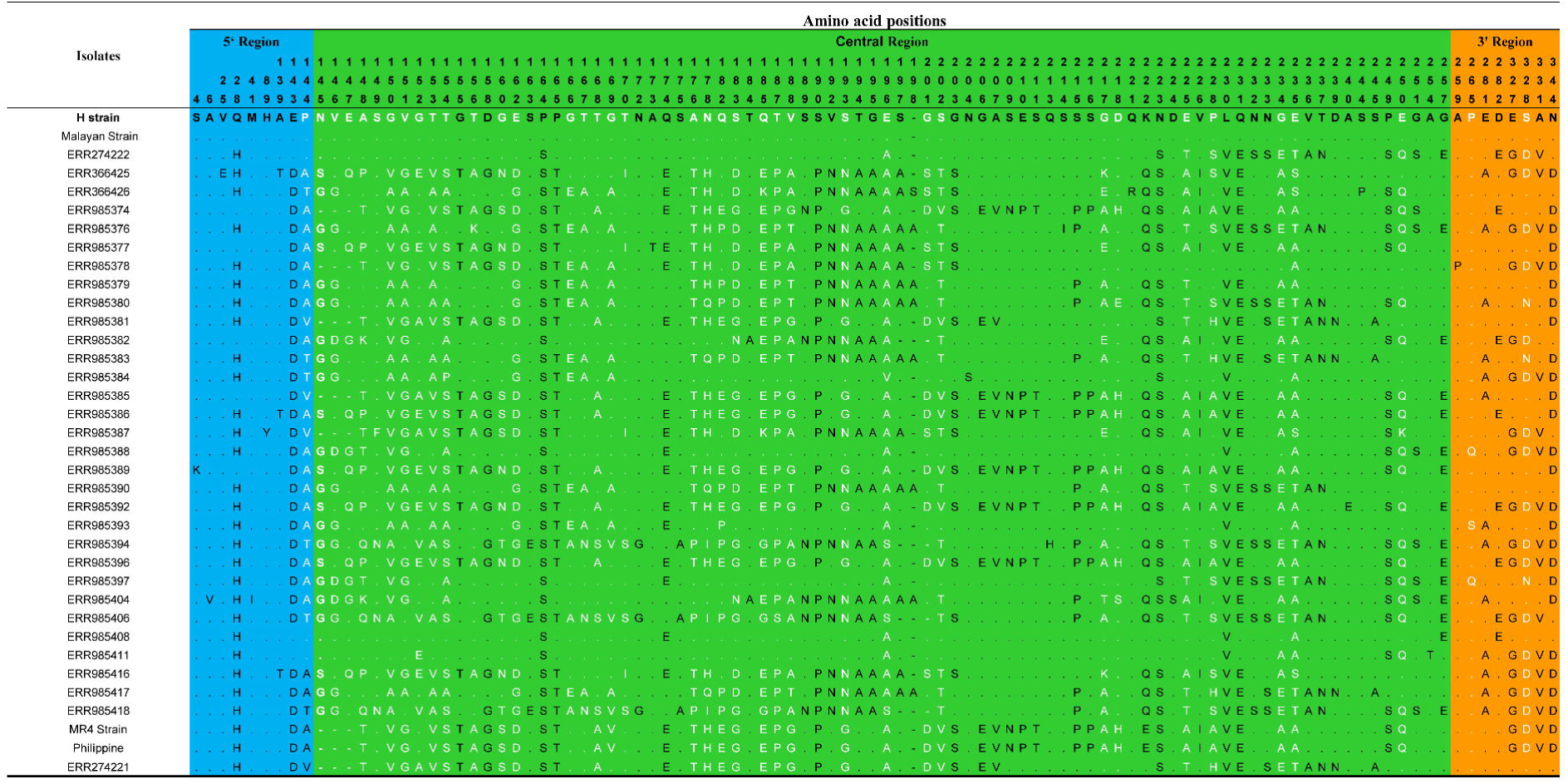
Amino acid polymorphism within 36 full-length PkMSP7D protein. The number above each amino acid denotes position of the amino acid based on the H-strain. Hyper-variable amino acids are highlighted in white. The blue, green and orange shaded regions represents the 5’ region, central region and the 3’ region of the PkMSP7D protein. The dots represents identical amino acids and dashes represents a deletion.

**Fig. 3.**
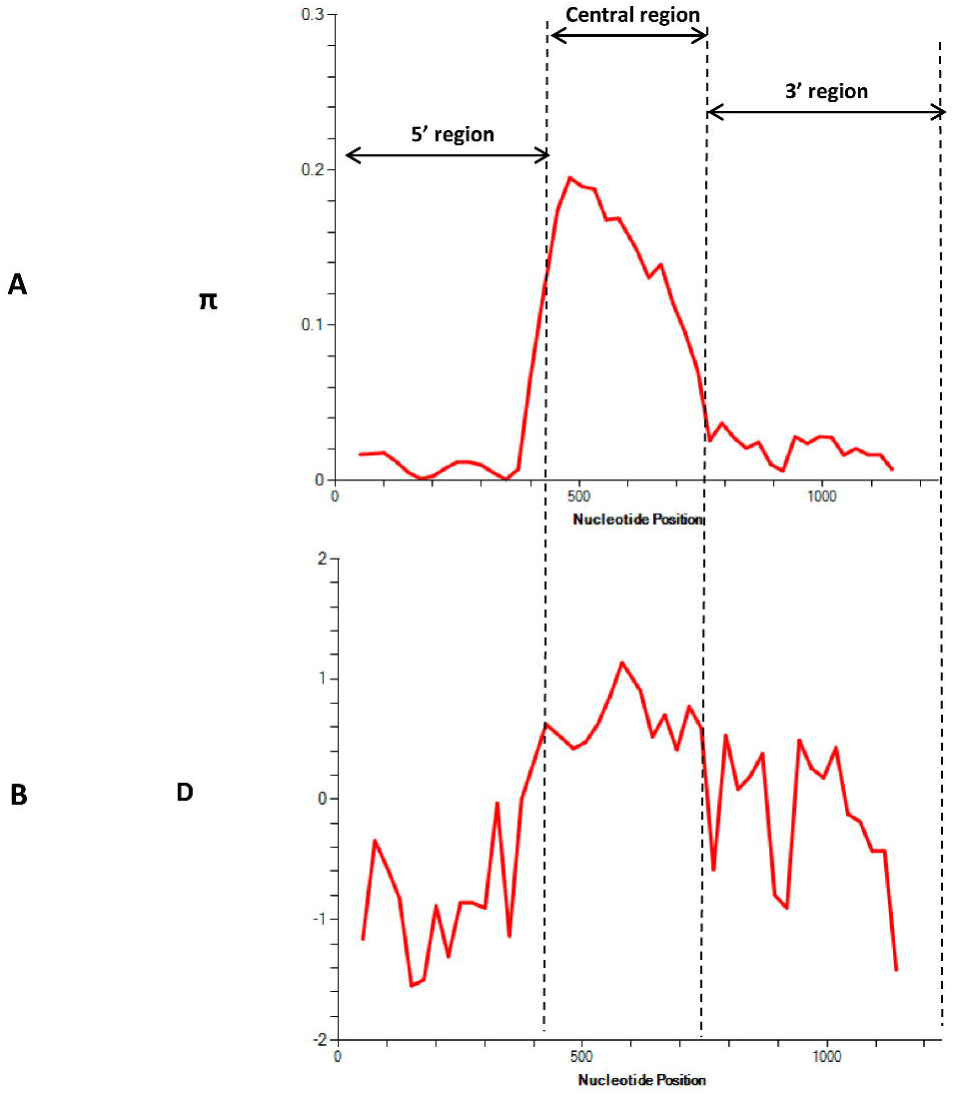
A Graphical representation of nucleotide diversity (π) within 36 full-length *pkmsp7D* genes from Malaysia. The *pkmsp7D* regions are marked as dashed line based on the diversity. (B) Graphical representation of Tajima’s D value across the *msp7D* gene.

### Region-wise analysis of diversity and natural selection

Based on the sequence alignment of *pkmsp7D* and the diversity pattern across the full-length gene (Fig. 3A), we defined three regions; 5’ region (from amino acid 1 to 144), central region (from amino acid 145 to 257) and the 3’ region (from amino acid 258 to 397). Nucleotide diversity at the 5’ and the 3’ regions were of similar levels (π = 0.010-0.017) (Table 1). The number of haplotypes in both the regions were 30 with similar levels of haplotype diversities (Hd = 0.992-0.990) (Table 1). Within the 5’ region, there were 28 SNPs (18 synonymous and 10 non-synonymous substitutions) and the 3’ region had 30 SNPs (22 synonymous and 8 non-synonymous substitutions) (Table 2).

The central region constituted the highest number of SNPs (151) leading to 32 haplotypes (Hd = 0.995) and the nucleotide diversity was very high (π = 0.149) compared to the other regions (Table 1). Of the 151 SNPs, 89 SNPs (16 synonymous and 73 non-synonymous substitutions) could be analysed using DnaSP software. Size variations within the *msp7D* genes were observed due insertion/deletion at two specific locations within the central domain; 6 nucleotides deletion (from 435-440nt) and 6-12 nucleotides insertion/deletion (from 590-601nt) in some of the isolates (Additional file 3). Twenty-three tri to tetra morphic nucleotides (hyper-variable SNPs) were observed within the central region (Additional file 3).

To determine whether natural selection contributes to the polymorphism in the 3’, 5’ and the central regions of *pkmsp7D* gene, the difference of (d_N_ - d_S_) and codon based Z test was evaluated independently for each region. The negative values at both the 5’ and 3’ region obtained indicated negative/ purifying selection (Table 1). The 3’ region was under strong negative selection pressure (dN-dS = 2.68, *P < 0.01*). Tajimas’D test, Fu and Li’s D* and F* test were all negative implying negative/purifying selection due to functional constrains at both these regions (Table 1). Negative selection was also evident as there were an excess of synonymous substitutions in both these regions (Table 2). Additionally, the robust MK test, which compares natural selection in the inter-species level, also showed significant values for both *P. vivax* and *P. cynomolgi* as outgroup sequences for the 3’ region (Table 3). Though MK did not yield statistically significant results for the 5’ region, the results were indicative of a negative selection.

The test results for natural selection for the highly diverse central region of *pkmsp7D* indicated the region was under the influence of strong positive/balancing selection. The difference of d_N_ - d_S_ = 1.84 (*P <*0.03) and Tajimas’ D, Fu and Li’s D* and F* values were also positive (Table 1). This was also evident due the presence of excess of non-synonymous substitutions within the region (nsSNPs = 73, sSNPs = 16) (Table 2). MK test, which also showed significant values when both *P. vivax* (NI = 2.49, *P <* 0.05) and *P. cynomolgi* (NI = 2.28, *P <* 0.05) were used as outgroup sequences for the central region (Table 3).

### Recombination and Linkage disequilibrium (LD) analysis

Analysis of recombination within the aligned *pkmsp7D* sequences estimated a minimum number of 29 recombination events (Rm) indicating intragenic recombination could be a factor adding to the extensive diversity at the central domain of the gene. LD analysis across the full-length *Pkmsp7D* genes showed the relationship between nucleotide distance and *R^2^* index and the decline in regression trace line indicated that intragenic recombination might exist within the *msp7D* genes in *P. knowlesi* (Fig 4).

**Fig. 4.**
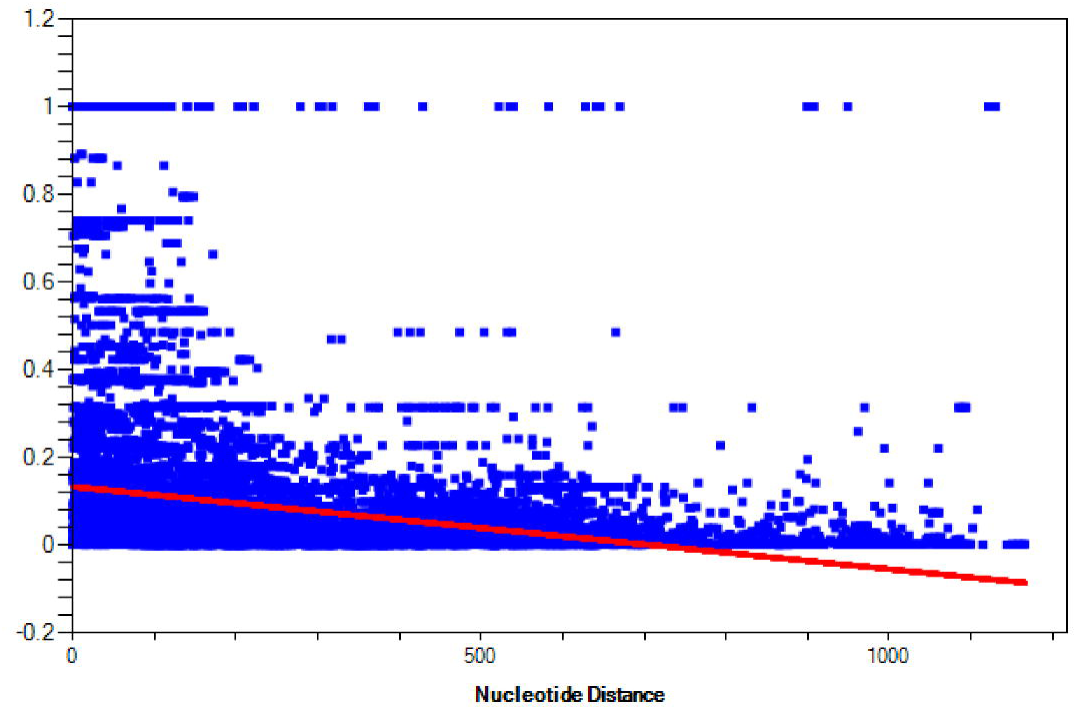
Linkage disequilibrium across the full-length *pkmsp7D* genes from Malaysia. Relationship between nucleotide distance and *R^2^* index and the decline in regression trace line indicated intragenic recombination within *P. knowlesi* isolates.

### Phylogenetic analysis

Phylogenetic analysis of the 37 full-length PkMSP7D amino acid sequences indicated that there was no geographical clustering of *knowlesi* sequences (Fig.4). The PkMSP7D sequences were found to be more closely related to *P. vivax* MSP7H compared to its paralogs in *P. knowlesi* and ortholog in *P. vivax* and *P. coatneyi* The H-strain and the Malayan strain clustered together along with sequences from Malaysian Borneo. The Philippine, MR4 and the Hackeri Strains formed a separate group along with Malaysian Borneo isolates. The distinct sub-populations observed in other invasion gene of *P. knowlesi* like MSP1P, DBPRII, NBPXA and TRAP was not observed in MSP7D [23, 25–27, 31, 49]. (Fig.5).

**Fig. 5.**
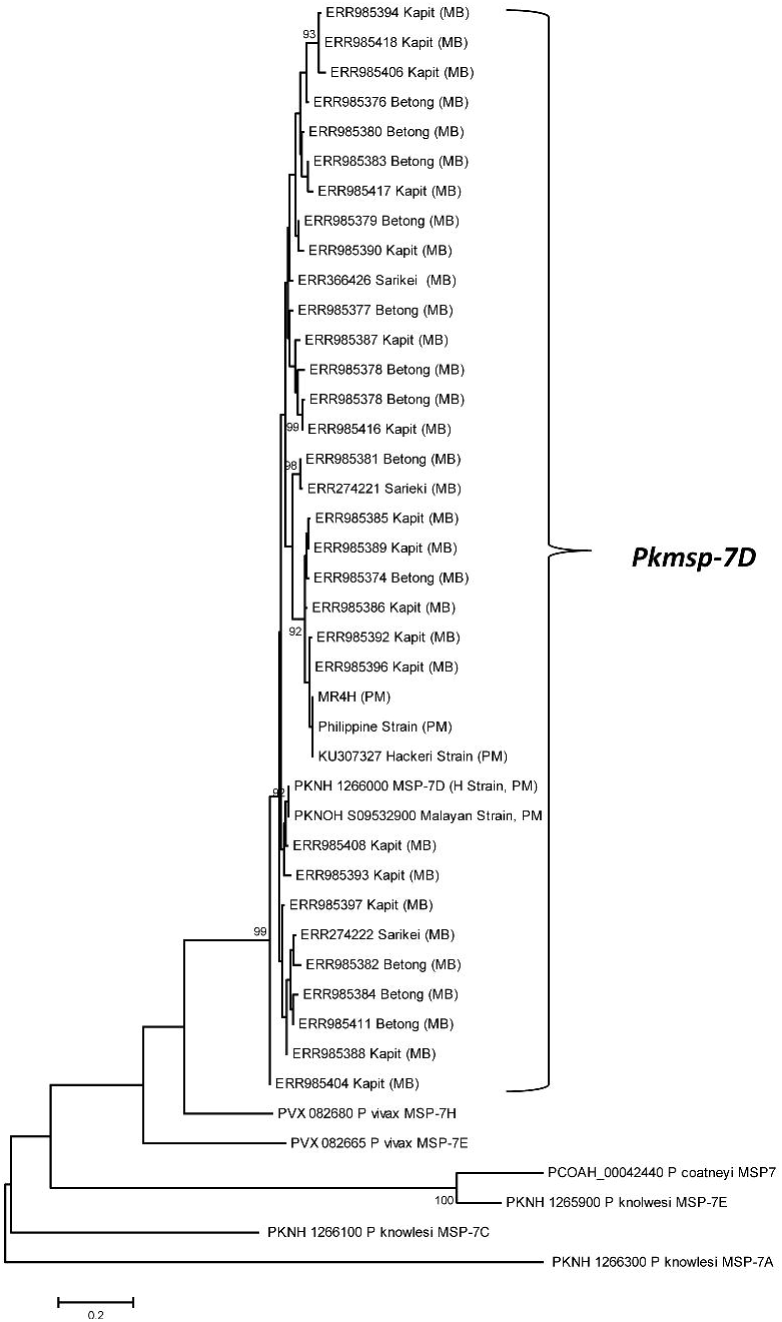
Phylogenetic relationship of deduced amino acid sequences of PkMSP7D from Malaysia and its paralog members PkMSP7A (PKNH_1266300), PkMSP7E (PKNH_1265900) and PkMSP7C (PKNH_1266100) and ortholog members of *Plasmodium vivax* MSP7H (PVX_082680) and *Plasmodium coatneyi* (PCOAH_00042440). The tree was constructed using Maximum Likelihood (ML) method based on Poisson correction model was used as described in MEGA 5.0 with 1,000 bootstrap replicates to test the robustness of the trees. MB and PM indicates samples from Malaysian Borneo and Peninsular Malaysia respectively

## Discussion

The merozoite surface protein (MSP7) gene family is a group of surface proteins which are involved in the initial interaction with MSP1 through a complex formation at the merozoite surface during the invasion process and orthologs and paralog members have been identified across *Plasmodium* species [37, 43]. In *P. falciparum*, MSP7 is expressed during schizont stages and undergoes two step proteolytic processing similar to MSP1 and disruption of the MSP7 gene leads to invasion impairment indicating the molecule is essential for the parasite [35]. Considering previous studies on MSP7 members in *P. vivax* (i.e. PvMSP7C, MSP7E, PvMSP7H and PvMSP7I) the central region were found to be under the influence of strong positive natural selection and extensive diversity indicating host immune pressure [38, 50]. Extensive functional as well as field based studies has been conducted on *P. falciparum* and *P. vivax* MSP7 gene family members, but no studies have been done on *P. knowlesi.* Thus, this study was conducted to genetically characterise the MSP7D genes from clinical isolates of Malaysia which is the closest ortholog of *P. vivax* MSP7H.

Sequence alignment of 36 full-length amino acid sequences of *pkmsp7D* genes from Malaysia showed that it shares approximately 60.4% sequence identity with its ortholog *Pvmsp7H* and the overall diversity across the full-length gene was higher than *P. vivax* [38]. Interestingly, the sequence identity within the PkMSP7 paralog member were low. Analysis of polymorphism across the full-length *pkmsp7D* gene indicated that only the central region acquired high number of polymorphism (SNPs =151) compared to the the 5’ and 3’ regions. This finding was similar to the diversity in the *msp7* members (C, E, H and I) in *P. vivax* [38, 50]. The genetic diversity at the central region of the gene was 14 fold higher (π = 0.1495) than the 5’ and 3’ region. This was due to mainly two reasons; firstly due to size variations in some of the clinical isolates in two specific locations within the central region and secondly due the high number of non-synonymous hyper-variable SNP sites (tri-morphic and tetra-morphic). The number of singletons (low frequency polymorphisms) in each of the regions were high leading to high number of haplotypes and haplotype diversity (Table 1). Similar hyper-variable sites were also observed in *P. vivax msp7* sequences from Thailand [50]. Tests for natural selection (using Taj D, Li and Fu’s D*, F*, dN-dS and MK test) at each region indicated that only the central region was under strong positive selection indicating immune evasion mechanism of parasite at this region, however, the 5’ and the 3’ prime regions are under functional constrains (purifying selection). These regions could potentially be the putative B cell epitope binding regions. This was probably due to the structural conformity attained by the protein during the invasion process and exposure of the central region to host immune system. Indeed, it remains to be proven through functional studies whether PkMSP7D forms a complex with PkMSP1 as observed in PfMSP1-MSP7 complex [34, 42]. A recent genetic and structural study on *P. vivax msp7E* sequences from Thailand predicted 4-6 α-helical domains at the 5’ and 3’ regions suggesting structural and functional constrains in these domains [50]. Sliding window analysis of Tajimas’D across the full-length *Pkmsp7D* identified high D values within the central region and a short region within the 3’ region. These regions with elevated D values were probable epitope binding regions.

Previous studies identified two sympatric sub-populations from Sarawak and geographical separation in samples from Peninsular Malaysia [24], however, phylogenetic analysis of the current study did not show any specific separation of the *P. knowlesi* MSP7D isolates. This is an interesting finding because the *pkmsp7D* sequences used in this study is from the same genomic data [24]. Bifurcation of trees, indicating dimorphism and strong negative/purifying selection on invasion genes like DBPαII (PkDBPαII) [31], PkNBPXa [26], PkAMA1 [30] have been reported from clinical samples. This probably indicates that *Pkmsp7D* is not influenced by geographical origin of the parasite and both the parasite sub-populations were under strong host immune pressure at the central region similar to the non-repeat region in *Pkcsp* from Malaysia [51]. We also found high recombination values and LD analysis indicated that intragenic recombination could play a crucial role in increasing diversity at the central region. These findings will be helpful for future designing functional binding assays to validate the molecule as a vaccine candidate.

## Conclusion

This study is the first to investigate genetic diversity and natural selection of *pkmsp7D* gene from clinical samples from Malaysia. High level of genetic diversity and strong positive selection pressure was observed in the central region of the *pkmsp7D* gene indicating exposure to host immune pressure. Absence of both dimorphism and geographical clustering within the parasite populations indicated uniform exposure of the central region to host immunity. Future studies should focus on functional characterization of *pkmsp7D* as a vaccine candidate.

## Abbreviations

MSP17: Merozoite Surface Protein 7 paralog
kDa: Kilodalton

## Declarations

### Acknowledgements

The authors would like to thank Dr. Syeda Wasfeea Wazid for helping in data retrieval and organizing the data for analysis.

### Ethics approval and consent to participate

Not applicable

### Consent for publication

Not applicable

### Availability of data and material

The datasets analysed during the current study were derived from the following public region resources: https://doi.org/10.1371/journal.pone.0121303 and https://doi.org/10.1073/pnas.1509534112

### Authors’ contributions

MAA and FSQ designed the study. MAA retrieved *Pkmsp7D* gene sequences from genomic databases, performed the genetic analysis. MAA and FQS wrote the manuscript. All authors read and approved the final version of the manuscript.

---

**Additional file 1 Table S1. Accession number of PkMSP7D sequences used in the study and their geographical origin**

**Additional file 2 Fig. S2. Signal peptide prediction by Signal IP server. Signal peptide was predicted in between amino acid positions 20 to 25**.

**Additional file 3 Nucleotide polymorphism within 36 full-length *pkmsp7D* sequences from Malaysia. SNPs with deletions are highlighted in red. Hyper-variable SNPs are highlighted in white. Blue, green and orange shaded regions indicate 5’ region, central region and 3’ region respectively**.

